# Evolution of a bistable genetic system in fluctuating and non-fluctuating environments

**DOI:** 10.1101/2024.01.22.576666

**Authors:** Rocío Fernández-Fernández, David R. Olivenza, Esther Weyer, Abhyudai Singh, Josep Casadesús, María Antonia Sánchez-Romero

**Affiliations:** Departamento de Microbiología y Parasitología, Facultad de Farmacia, Universidad de Sevilla, Calle Profesor García González 2, 41012 Sevilla, Spain; Departamento de Genética, Facultad de Biología, Universidad de Sevilla, Avenida Reina Mercedes 6, 41012 Sevilla, Spain; Departments of Electrical and Computer Engineering, Biomedical Engineering, Center for Bioinformatics and Computational Biology, University of Delaware, Newark, DE 19716, U.S.A

## Abstract

Epigenetic mechanisms can generate bacterial lineages capable of spontaneously switching between distinct phenotypes. Currently, mathematical models and simulations propose epigenetic switches as a mechanism of adaptation to deal with fluctuating environments. However, bacterial evolution experiments for testing these predictions are lacking. Here, we exploit an epigenetic switch in *Salmonella enterica,* the *opvAB* operon, to show clear evidence that OpvAB bistability persists in changing environments but not in stable conditions. Epigenetic control of transcription in the *opvAB* operon produces OpvAB^OFF^ (phage-sensitive) and OpvAB^ON^ (phage-resistant) cells in a reversible manner and may be interpreted as an example of bet-hedging to preadapt *Salmonella* populations to the encounter with phages. Our experimental observations and computational simulations illustrate the adaptive value of epigenetic variation as evolutionary strategy for mutation avoidance in fluctuating environments. In addition, our study provides experimental support to game theory models predicting that phenotypic heterogeneity is advantageous in changing and unpredictable environments.

---

Microbial populations are faced with multiple challenges in natural environments and have evolved and adapted to survive in a wide range of natural conditions ^1,2^. An especially severe threat is the encounter with bacteriophages. Considering the abundance and ubiquity of these biological entities, one may thus understand the adaptive value of mechanisms that protect from bacteriophage infection ^3,4^. Indeed, bacteria have evolved a bewildering array of mechanisms of defence against bacteriophages ^5^. One such mechanism is the reversible (“phase-variable”) formation of bacterial subpopulations^6^ able to survive in the presence of a virulent phage ^7–12^. This phase variable system is an epigenetic switch that allows the existence of multiple stable phenotypic states in a genetically identical population. The occurrence of phase-variable phage resistance in unrelated bacterial species and the diversity of mechanisms involved suggest independent evolution, which in turn may be indicative of adaptive value.

In *Salmonella enterica*, OpvA and OpvB proteins control O-antigen chain length in the lipopolysaccharide (LPS) ^10^, offering resistance to bacteriophages that use the O-antigen as receptor ^11^. The *opvAB* operon, regulated by the OxyR transcription factor and DNA methylation, governs OpvA and OpvB expression, leading to bistable switching between ON and OFF states ^13^. The bistable switch is lopsided towards the OFF state, and, although mostly cells are in the OFF state, the encounter with a phage selects for the survival of OpvAB^ON^ subpopulation (Fig. 1a). As soon as phage challenge ceases, phase variation rebuilds a mixed population made of OpvAB^OFF^ and OpvAB^ON^ lineages ^10,11^. Because OpvAB^ON^ cells are impaired for interaction with the animal host, phase variation of O-antigen length can be viewed as a tradeoff between pathogenic capacity and bacteriophage resistance. The OpvAB^ON^ subpopulation of *Salmonella* preadapts to phage challenge survival, albeit at the expense of reduced virulence ^11,14^.

**Figure 1.**
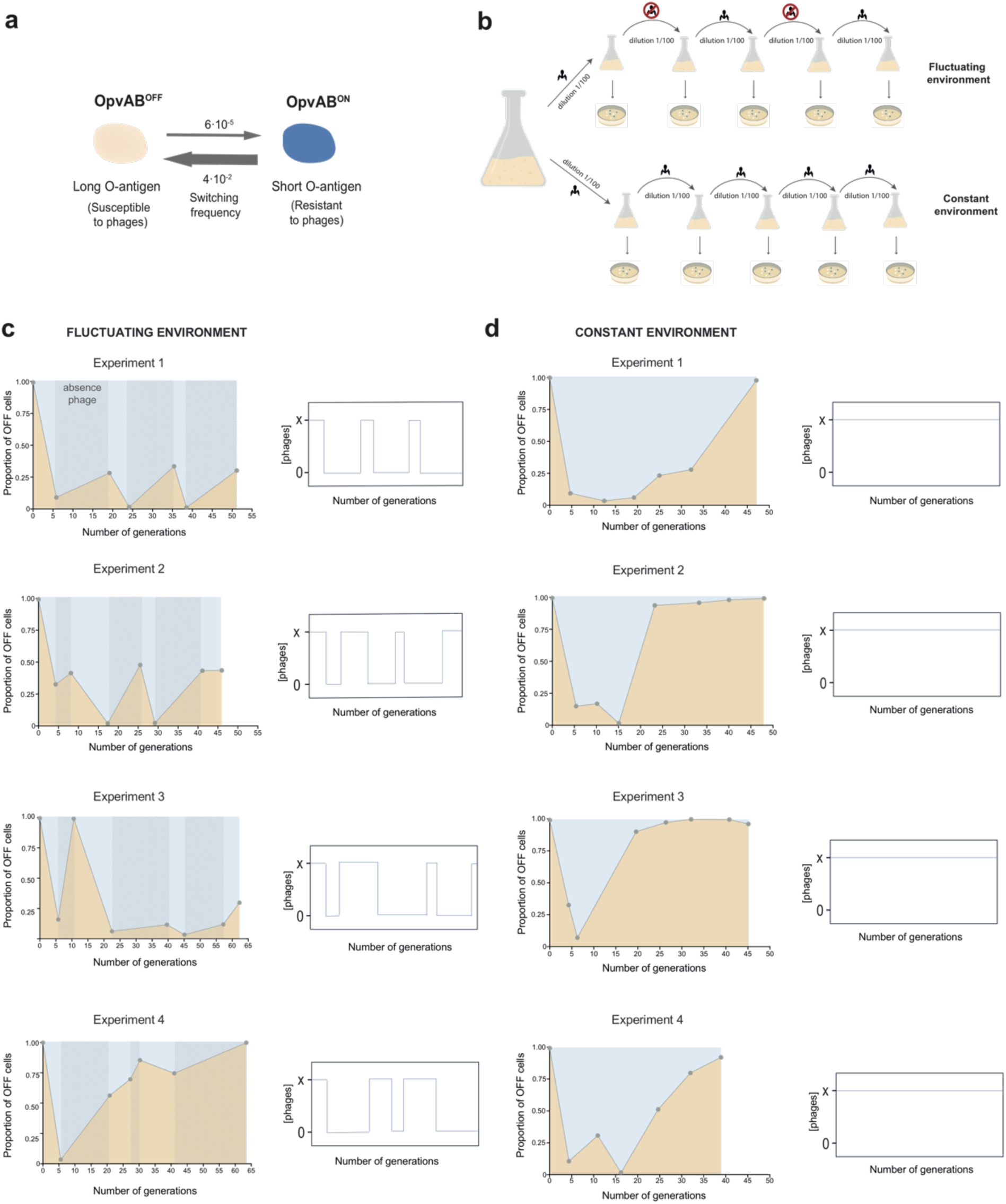
Evolution of the bistable *opvAB* genetic system in fluctuating *versus* constant environments. **a.** *Salmonella enterica opvAB* expression undergoes phase variation and produces a lineage of OpvAB^OFF^ cells with long O-antigens in the lipopolysaccharide that are susceptible to phages, and a lineage of OpvAB^ON^ cells with shorter O-antigen chains resistant to phages. Switching from OFF to ON states occurs at lower frequencies than switching from ON to OFF. The frequency of OFF to ON transition was estimated to be 6·10^-5^ per cell and generation. The ON to OFF switching rate was around 1,000-fold higher: 4·10^-2^ per cell and generation. Proportion of OFF and ON cells in the bacterial population is around 99.8% and 0.2%, respectively (Cota et al, 2012). **b.** Scheme of the laboratory evolution experiment to simulate fluctuating and constant environments. For fluctuating environment, presence and absence of bacteriophages were used to simulate a changing environment. A dilution of an overnight culture of 14028 *opvAB::lacZ* strain was performed in presence of phage and plated LB supplemented with X-gal agar with phages. Phage removal protocol was carried out to remove phage number and cease the challenge. Six alternating cycles of phages presence and absence were performed. For constant environment, cultures were achieved by diluting an overnight culture of 14028 *opvAB::lacZ* strain in presence of 9NA lysate, plating on LB supplemented with X-gal plates and phages each 24 hours. **c and d.** Proportion of OpvAB^OFF^ colonies of the strain *opvAB::lacZ* growing in fluctuating (c) and continuous presence of bacteriophage (d). Four independent laboratory evolution experiments were performed. Yellow shades area under the curve represents the proportion of OFF cells and blue shades area above the curve shows the proportion of ON cells in the bacterial culture. Dotted grey shaded areas indicate the absence of phages in fluctuating environment. Simplified diagrams clarify the presence and absence of phages over generations.

Phenotypic switching has been proposed as a mechanism of bet-hedging to deal with fluctuating environments ^15–21^. Specifically, formation of OpvAB^OFF^ and OpvAB^ON^ lineages may be seen as an example of bet-hedging ^22^, which allows *Salmonella* populations to prepare themselves for potential encounters with phages ^11^. Natural selection of phenotypic heterogeneity is a controversial notion in classical Darwinism as it involves group selection, which has been traditionally considered a weak evolutionary force ^23^, a criticism especially severe when formation of lineages is interpreted as a case of bet hedging ^24^. This view is however countered by game theory models ^25–27^. Formation of distinct types of cells preadapted to different environments is also in agreement with theoretical models by Richard Levins showing that polymorphism is advantageous in changing environments ^28,29^. Even though Levins’s studies were intended for genetic polymorphism, their conclusions could potentially be extended to any source of variation, including formation of epigenetic lineages. Natural selection operates on phenotypes, not on genotypes ^30,31^.

To test Levins’s prediction that polymorphism increases the fitness of biological populations in changing environments but not in stable ones, conducting bacterial evolution experiments combined with mathematical modelling approaches is necessary. This analysis will allow to evaluate how microbial lineages can respond to variable and heterogeneous environments both theoretically and practically. Here, we exploit the well-known epigenetic switch based on *opvAB* operon, which generates bacterial subpopulations differing in specific phenotypic traits, to shed light on the factors by which epigenetic mechanisms contribute to *Salmonella*’s ability to adapt and persist in unpredictable natural environments.

## Results

### OpvAB subpopulation proportions vary depending on the fluctuations of the environmental conditions

We exploited the *opvAB* gene expression system to investigate the variation of OFF and ON phenotypic states in fluctuating and non-fluctuating environments (Fig. 1a,1b). We used a *Salmonella enterica* strain harbouring an *opvAB::lacZ* fusion (SV8011) to conduct four independent laboratory evolution experiments in different environmental conditions (Fig. 1b). The strain SV8011 was grown with and without bacteriophages, and white (OpvAB^OFF^) and blue (OpvAB^ON^) colonies were counted. In fluctuating environment, alternate growth cycles with and without bacteriophages were repeated and colony counts were recorded (Fig. 1b,1c). Phage-free medium was performed after removal of phage by treatment with purified LPS. In constant environment, SV8011 was growth in presence of phages, and the bacteriophage removal step was omitted (Fig. 1b,1d). The results in these experiments (Fig. 1c,1d) reveal distinct patterns. In alternating phage presence (Fig. 1c), the proportion of OpvAB^OFF^ colonies decrease in phage presence, rebounding when absent. Conversely, the proportion of OpvAB^ON^ colonies exhibit an opposite trend. In continuous phage exposure (Fig. 1d), the proportion of OpvAB^ON^ initially increases, but unexpectedly decreases with distinct speed and kinetics at each experiment after 5-10 generations, leading to near-extinction. OpvAB^OFF^ colonies then took over the culture after 40-50 generations.

The numerical data were slightly different in each experimental replicate, but the overall phenomenon remained the same: OpvAB^ON^ and OpvAB^OFF^ colonies are present throughout the experiment carried under fluctuating selection, whereas OpvAB^OFF^ colonies overtake the culture growing under continuous selection with a virulent phage. Importantly, this laboratory evolution experiment supports Levins’ prediction ^28,29^: bistability is favoured by maintaining different subpopulations in fluctuating environments. In contrast, when the environmental conditions are constant, phenotypic heterogeneity is lost and a monostable cell populational distribution becomes established.

### OpvAB^OFF^ isolates from cultures grown in the continuous presence of phage are predominantly mutants

Since OpvAB^ON^ colonies are made of phage-resistant cells (Fig. 1a), survival upon bacteriophage infection explains their predominance over OpvAB^OFF^ colonies under fluctuating conditions (Fig.1c). In contrast, the number of OpvAB^ON^ colonies progressively decreases under continuous selection, dropping to near-extinction; OpvAB^OFF^ colonies thus become the predominant class (Fig. 1d). Because OpvAB^OFF^ cells have long O-antigen chains, the survival of OpvAB^OFF^ colonies from the constant-environment evolutionary experiment in the presence of phage must be made possible by mechanism(s) unrelated to *opvAB* phase variation. One potential mechanism could be mutation. To investigate if mutational genetic adaptation is responsible for the survival of OpvAB^OFF^ cells in the presence of phage, we subjected twenty phage-resistant OpvAB^OFF^ isolates to whole DNA genome sequencing. All of them showed mutations in their genomes (356 different mutations were detected). Two thirds mapped to coding sequences (97.5% in known genes and 2.5% in genes annotated as hypothetical proteins) and one third to intergenic regions. As a result, 72 genes encoding proteins with known function were affected (Supplementary Table 1, Fig. 2). More than 50% (38 out of 72 genes) of these genes were altered in more than one phage-resistant isolate (Fig. 2a) and nearly 87% (33 out of 38 genes) presented more than one mutation in the same gene, predominantly substitutions (Fig. 2b,2c). All these observations suggest that these mutations were not randomly distributed. The analysis of mutated genes reveals a high prevalence of genes related with phage infection and subsequent bacterial lysis (Fig. 2d). In addition, genes associated with virulence and cell permeability were notoriously affected. Other biological processes compromised in phage-resistant isolates were related to translation, carbohydrate metabolism and cell wall and membrane biogenesis (Fig. 2d). Most of these processes have been also reported to be altered in phage-resistant isolates from other Gram-negative bacteria, such as *Escherichia coli* or *Vibrio alginolyticus* ^32–34^, suggesting their robust implication in mechanisms of phage resistance.

**Figure 2.**
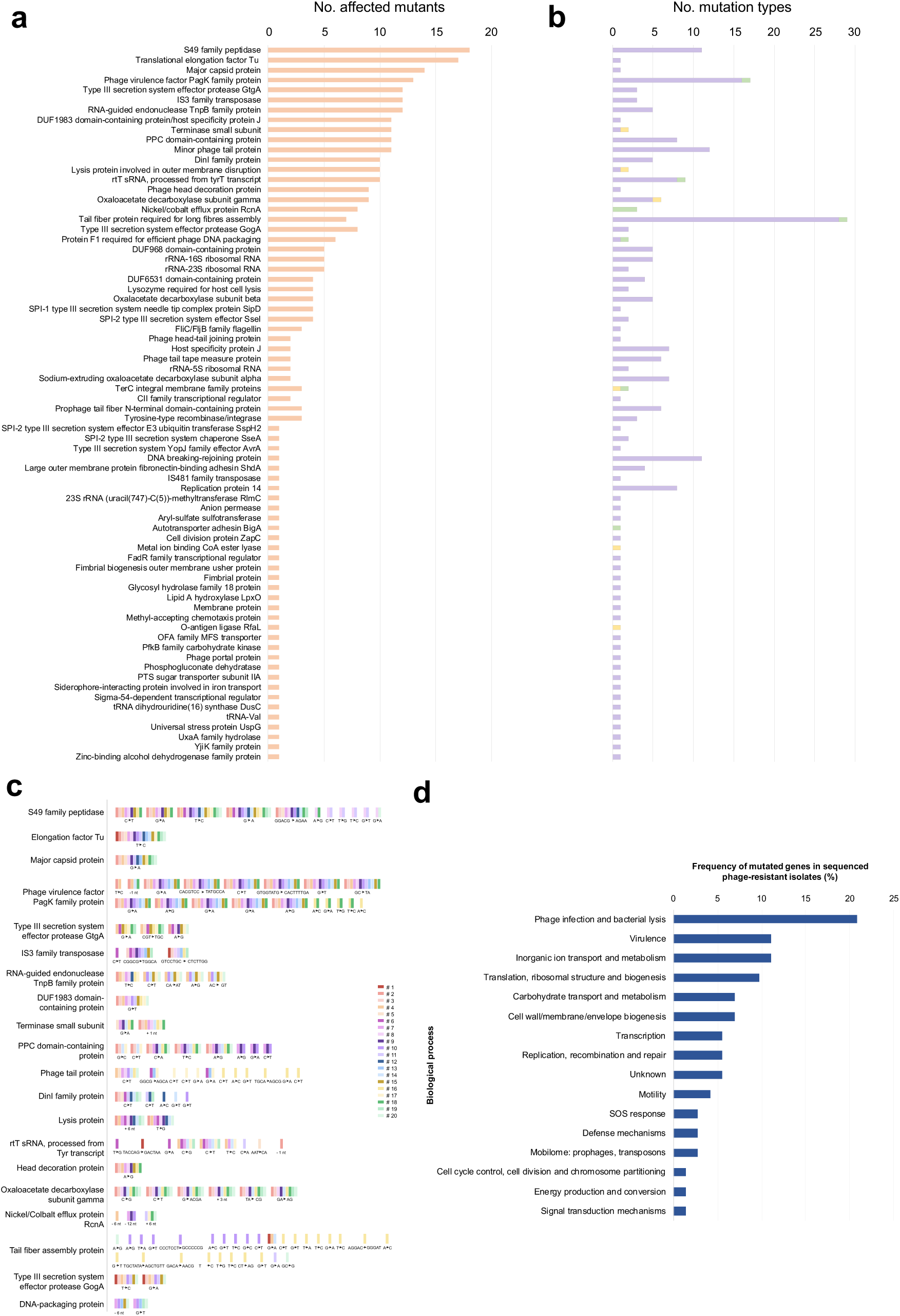
Description of the mutations detected in OpvAB^OFF^ phage-resistant isolates. **a and b.** Identification of altered genes found in the DNA sequencing of OpvAB^OFF^ phage-resistant isolates. For each affected function, predominancy in phage-resistant isolates (out of 20) (a, orange) and type of mutations (b, substitutions (violet), insertions (yellow) and deletions (green)) found in each gene are represented. **c.** Analysis of types of most common mutations found in the twenty OpvAB^OFF^ phage resistant isolates. Altered functions shown are shared by at least 30% of the mutants (6 out 20). Types of mutations are identified. Colors identified the number of phage-resistant isolates. **d.** Clusters of biological process classification and distribution (frequency/incidence) of mutated genes in sequenced phage-resistant isolates.

In summary, DNA sequencing of OpvAB^OFF^ isolates from cultures grown in the continuous presence of bacteriophages demonstrated the existence of mutations related to phage resistance, supporting the notion that prolonged exposure to phages influences the genetic adaptation and favours a more stable phenotype over bistability.

### Mathematical modelling of the bistable *opvAB* epigenetic system

To further explore the evolution of the bistable gene expression in *opvAB* system under diverse environmental conditions, firstly, we constructed a mathematical model for simulating the *opvAB* operon subjected to epigenetic control by DNA methylation. Secondly, we evaluated how the rates between *Salmonella* OpvAB^OFF^ and OpvAB^ON^ cell subpopulations evolved in contact with bacteriophages using different model assumptions.

Interaction dynamics in *Salmonella* cell population containing the *opvAB* operon in presence of phage is illustrated in Fig. 3a. To start with, we simulated the evolution of a bacterial population in an environment where phages were always present (Fig. 3b,3c) or where absence and presence of phages fluctuated, by sequentially removing and re-introducing phages over several intervals (Fig. 3d,3e). The predicted fraction of ON and OFF cell states was plotted as a function of time (Fig. 3b,3d).

**Figure 3.**
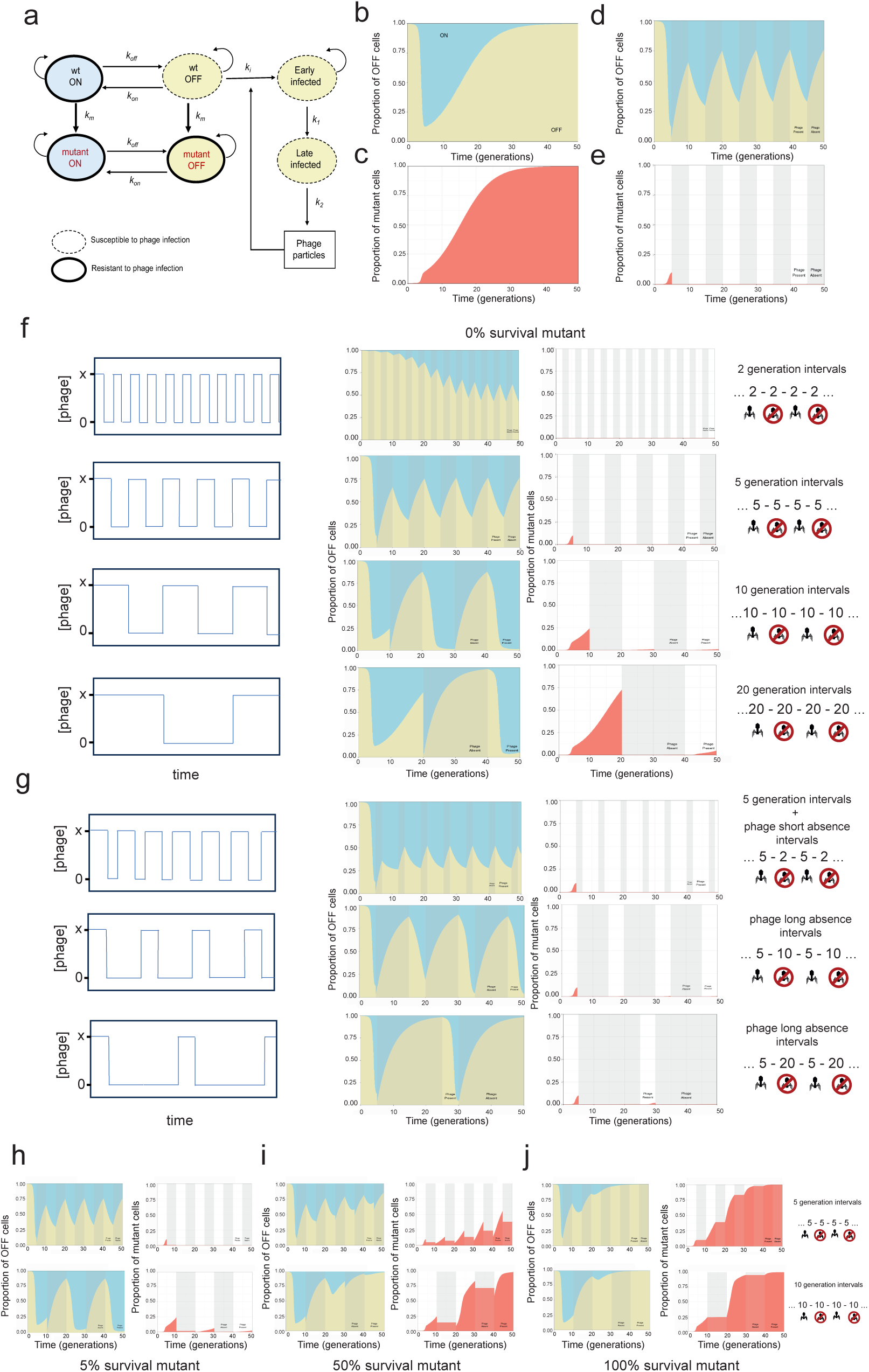
Modelling the bistable *opvAB* system in diverse environmental conditions. **a.** Flow chart of populations and interaction dynamics used for model. Uninfected cells replicate and switch between OFF and ON states. Wild-type cells permanently mutate and become resistant to the bacteriophage particles even in the OFF state. Susceptible OFF cells are infected by bacteriophage particles and continue to replicate until they become late infected and release new bacteriophage particles by bursting. **b and d.** Proportion of OpvAB^OFF^ cells predicted by the model with best fitting parameters from the constant (b) and fluctuating (d) environments. Predicted proportion of cells in the ON state was calculated by 1-proportion of OFF cells. **c and e.** Proportion of mutant cells predicted by the model using the best fitting parameters from the constant (c) and fluctuating (e) environments. **f.** Proportion of OpvAB^OFF^ cells and proportion of mutant cells predicted by the model using different intervals of phage absence and presence in the simulations from the fluctuating environment, assuming 0% mutant survival. **g.** Proportion of OpvAB^OFF^cells predicted by the model using different intervals of phage absence and presence in the simulations from the fluctuating environment, assuming 0% mutant survival. In these simulations is assumed 5 generation intervals in presence of phage but different absence interval times. **h-j.** Proportion of OpvAB^OFF^ cells and proportion of mutant cells predicted by the model using 5 and 10 generation intervals in the simulations from the fluctuating environment, assuming 5%, 50% and 100% mutant survival (h, i and j, respectively). Yellow shades area under the curve represents the proportion of OFF cells and blue shades area above the curve shows the proportion of ON cells in the bacterial culture along time. Red shades area under the curve represents the proportion of mutant cells over time. Dotted grey shaded areas indicate the absence of phages in fluctuating environments.

In the constant presence of bacteriophage, the proportion of OFF cells was predicted from the initial concentrations using equation 9 (Supplementary Fig. 1). The model is able to match the initial rapid decrease of the proportion of OFF cells after the introduction of bacteriophages and the subsequent gradual recovery to the initially observed proportions of OFF/ON subpopulations (majority of OFF cells), as shown experimentally (Fig. 3b and Supplementary Fig. 2). When the proportion of mutants was modelled, the mutant proportion shows a short delay before a rapid increase approaching 1 (Fig. 3c). This trend correlated with the previously observed experimental data, which showed the prevalence of OFF mutant cells in a constant environment. Thus, the model reproduces experimental data demonstrating that constant environment conditions favour the selection of mutations and decrease phenotypic diversity (Figs. 1d,3b,3c and Fig. Supplementary 2).

For the fluctuating environment, the model was roughly able to predict the shifting behavior of the OFF and ON proportions. Expectedly, the simulation illustrates the increase in the proportion of OFF cells as bacteriophage particles were removed and the upsurge in the ON proportion as phage particles were re-introduced (Fig. 3d). However, the prediction of population proportions in fluctuating conditions oscillated close to 0.5 values. This slightly differs from our previous experimental observations where the amplitude of oscillations varies between experiments (Fig. 1c). This discrepancy could be due to assuming cyclical fluctuations in intervals of 5 generations in presence and absence of phages for modelling the fluctuating environment. More complexities at the cycles of phages exposure in individual experiments could explain this difference. These points will be discussed later.

As a consequence of the procedure for bacteriophage removal, the number of generations (which depend on initial and final cell number in the culture at the end of the phage-challenge step) during each cycle of phage presence and absence varies across different experiments (Fig. 1c, shaded intervals). We ran several simulations changing the number of generations occurring in the presence and/or absence of phages (Figs. 3f,3g). For each scenario, OFF and ON subpopulations size fluctuates based on the point of occurrence of phages and the duration spent recovering in culture without phage. In general, the fluctuations in the proportions of ON and OFF population (amplitude of the oscillations) became much larger as frequency decreased. Therefore, in fast-changing environments, the population evolved to a bistable phenotype (Fig. 3f Top), whereas in slow-changing environments the tendency is for one phenotype to be predominant (Fig. 3f Bottom). When we conducted simulations with asymmetric cycles (phage-free intervals longer or shorter than intervals with phage presence), we observed that the amplitude of oscillations seems to correlate with the length of the interval without phage (Fig. 3g). The longer the culture is free of phages, the greater the amplitude of oscillations, and the higher the probability that one subpopulation dominates the culture for an extended period. These phenomena could be similarly observed in each of the replicates undergoing fluctuating environments (Fig. 1c). Hence, as the simulation indicates and our experimental data show, the period of environmental changes can modulate the OFF and ON subpopulations proportions generated by an epigenetic switching system.

As previously described, mutant cells become predominant under a constant phage pressure, consequently, genetic adaptation appears to be a better strategy in constant environments. Surprisingly, we found a similar behavior in the experiment 4 grown in fluctuating environments: the OFF cells take over the culture after 30-35 generations leading to the establishment of a single subpopulation (Fig. 1c). We wondered how mutant survival affects the behavior on OFF/ON proportions predicted by the model in changing environments. The initial assumption in our model was that no mutants survived at the dilution events, preventing the recovery of the initial OFF/ON proportions. In fluctuating environments, the jump in proportions that occurs after the first interval is due to the assumed loss of the mutant population (Fig. 3d). At the end of the first interval (5 generations in the presence of phages), the proportion of mutants is about 10% (Fig. 3e), so a gap occurs when the mutant concentration is set to 0 for the next initial conditions. When we considered 10 and 20 generation intervals, the proportion of mutants at the end of the initial interval was close to 25% and 75%, respectively (Fig. 3f), causing a much larger gap in the predicted proportions (Fig. 3f). Because total extinction is a strong assumption, the same interval scenarios were also tested with 5%, 50%, and 100% mutant survival rates (Figs. 3h-3j). When a survival rate of 5% was tested, no significative difference compared to the 0% survival was found (Fig. 3f,3h); however, a 50% mutant survival rate showed distinctive pattern of fluctuations in the different interval scenarios and the ON/OFF proportions noticeably began to approach the steady state much more rapidly (Fig. 3i). Because there was partial survival, the gaps became smaller at the end of the first interval, but then greater at the subsequent dilutions where the mutant population was higher than in the 0% survival cases. The increase in the mutant proportion during the intervals of phage presence is also more noticeable. It is worth highlighting that unlike what happens with the predicted OFF/ON proportions in 10 generations intervals and 0% mutant survival (Fig. 4f) where the ON subpopulation takes over the culture during phage presence intervals, when a 50% mutant survival is tested, the OFF subpopulation becomes predominant after 40-50 generations (Fig. 4i). A 100% survival mutant rate produced very similar results to the constant environment case (Fig. 4j), and OpvAB^OFF^ subpopulation takes over the culture more rapidly. These model parameters would explain the results obtained for the experimental replicate number 4 grown in fluctuating conditions (Fig. 1c).

**Figure 4.**
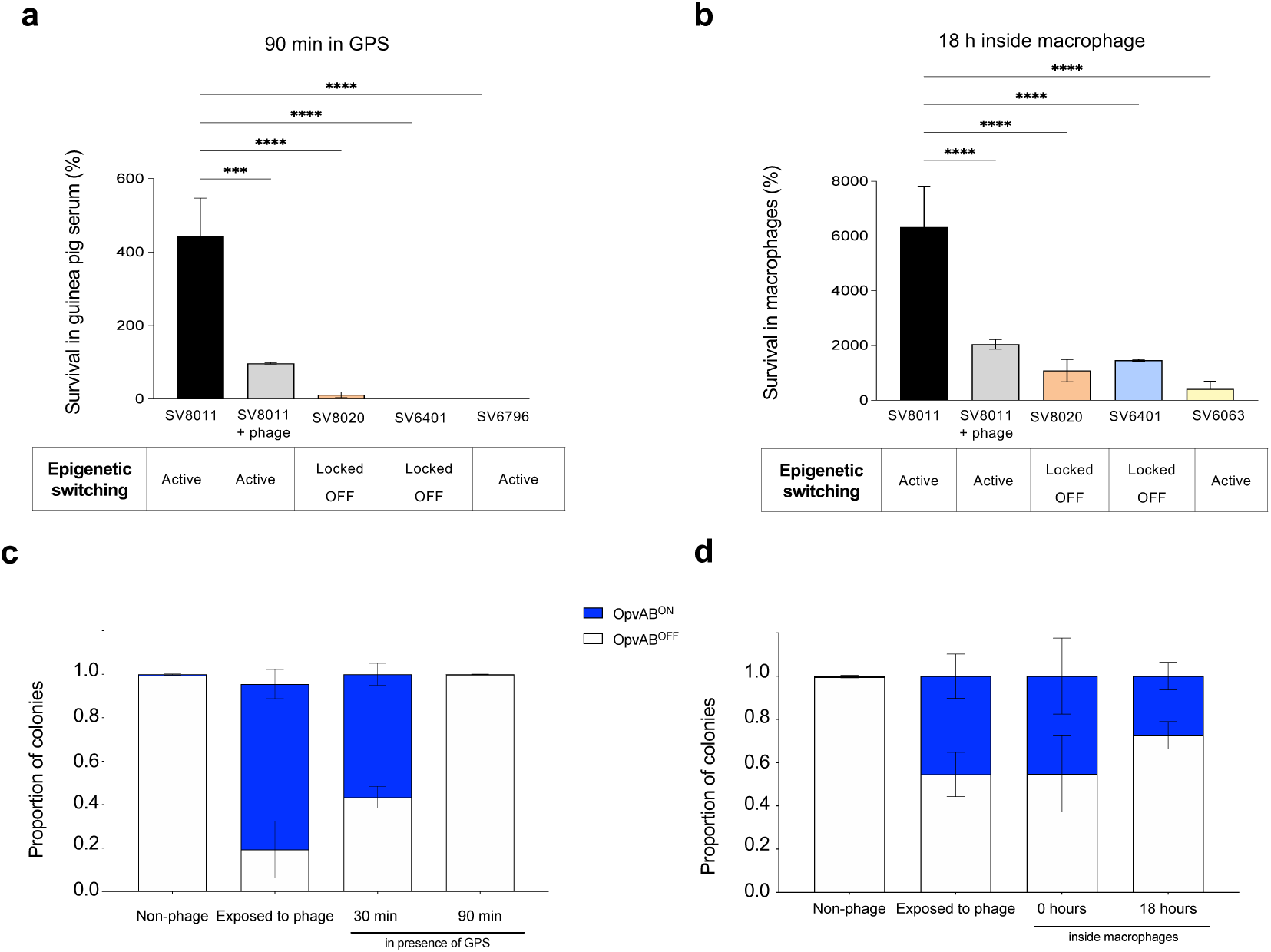
Role of reversible switching of *opvAB* expression for adaptation to diverse selective pressures. **a.** Survival rate in presence of 30% of guinea pig serum (GPS) for 90 minutes. Survival rate is relative to 0 minutes in response to GPS. **b.** Intramacrophage proliferation rate for 18 hours inside J774 murine macrophages cell line. Proliferation rate is relative to 0 minutes in contact with macrophages. **c.** Dynamic analysis of proportion of OpvAB^ON^ and OpvAB^OFF^ cells in presence of GPS that have previously been in contact with phage. Proportions were investigated in a wild-type strain (SV8011) before adding bacteriophage, after 10 generations with 9NA phage, immediately after adding 30% GPS and after 90 minutes of incubation in presence of GPS. **d.** Dynamic analysis of proportion of OpvAB^ON^ and OpvAB^OFF^ cells inside macrophages of the wild-type strain (SV8011) that had been in contact with bacteriophage before invasion of macrophages. Proportions shown correspond to the wild-type strain before adding bacteriophage, after 10 generations in presence of 9NA phage, 30 minutes inside macrophages and after 18 hours of proliferation in macrophages. Strains used for these studies were the following: SV8011, as wild-type strain harboring an *opvAB::lacZ* fusion, containing active epigenetic switching, that is, the majority of OpvAB^OFF^ cells and minority of OpvAB^ON^ cells (around 0.2%); SV8011 + phage is a culture that was previously in contact with 9NA phage for 10 generations, containing the majority of OpvAB^ON^ cells and minority of OpvAB^OFF^ phage-resistant cells, probably as mutants; SV8020 is a strain containing an *opvAB* deletion and an altered LPS profile, which exhibited phage resistance and bacterial cultures only show OFF phenotype; SV6401, a strain that remains blocked in the OpvAB^ON^ state and, thus, bacterial population only display ON cell phenotype. SV8020 and SV6401 strains present permanent lipopolysaccharide modifications and are phage-resistant. SV6796 strain is 14028Δ*wzz_ST_* Δ*wzz_fepE_* and is sensitive to serum. SV6063 is avirulent strain which has a deletion of *SPI-2* genes that makes it uncapable of proliferating inside macrophages.The experiment was performed in triplicate and averages and standard deviations are shown. Statistical indications: ns, not significantly different, **** P < 0.0001, *** P < 0.001, ** P < 0,01, * P < 0.05.

In summary, our model accurately predicts how the fluctuations in the presence/absence of phages and the survival rate of mutants significantly affect the structure of bacterial cell populations.

### Response of epigenetic switching for *opvAB* expression to multiple selective pressures

Microorganisms face the challenge of adapting to multiple selection pressures. In natural environments, where the presence of a selective pressure eliminated the mutants, the maintenance of the ON/OFF populations would offer the higher evolutionary process. The expression of the *opvAB* operon reduces the length of the O-antigen and, consequently, the absorption of bacteriophages, becoming resistant to these phages. Notably, the acquisition of phage-resistance by phase variation of O-antigen chain length involves an expensive payoff for a pathogen as *Salmonella*: *opvAB* expression impairs serum-resistance and reduces proliferation within macrophages ^10,11^. Thus, we hypothesized that mutants resistant to phages would experience a permanent reduction in virulence; on the contrary, the transient phase variation of *opvAB* gene expression would offer bacterial populations the versatility needed to confront a variety of adverse agents, such as phages and their hosts. To demonstrate this, we examined survival in guinea pig serum and intramacrophage replication in cultures containing cells that could reversibly switch (SV8020 strain) or be permanently locked in the ON (SV6401 strain) or OFF states (SV8020 strain, OpvAB^OFF^ lineage phage-resistant with altered LPS profile) (Fig. 4). Survival in guinea pig serum (GPS) was analyzed for 90 minutes (Fig. 4a) and the rate of intramacrophage replication was measured after 18 hours of infection in J774 mouse macrophages (Fig. 4b). Our results showed an increase of killing by serum and an impaired intramacrophage proliferation of those strains in which phase variation mechanism is abolished (SV6401 and SV8020 strains) compared to wild type (SV8011). Interestingly, when SV8011 cultures that were previously in contact with phage (thus containing the majority of OpvAB^ON^ cells but with active reversible switching) were analyzed, the serum-resistance and the ability to proliferate within macrophages increased compared to the strains with inactive *opvAB* phase variation (Fig. 4a and 4b). These observations support our hypothesis that the reversible switching of *opvAB* gene expression may be required for adaptation to changing environments. To confirm this hypothesis, we exposed wild-type cultures to bacteriophages for 10 generations prior to monitoring the proportion of OpvAB^ON^ and OpvAB^OFF^ subpopulations in the presence of serum (Fig. 4c) or before growth within macrophages (Fig. 4d). Highly dynamic OFF/ON proportions and drastic changes were observed: the initial proportion of OpvAB^ON^ cells was very small (0.2%) in the absence of phage, but, as soon as bacteriophages were added, the OpvAB^ON^ proportion increased rapidly, reaching values between 45-70%. However, after 90 minutes of serum treatment or 18 hours of intramacrophages replication, the OFF/ON proportions returned to the initial pre-phages levels and bacterial cultures were again enriched in OpvAB^OFF^ cells (serum-resistant and increased intramacrophage proliferation). Accordingly, survival in response to a new additional selective pressure (exposure to serum or macrophages) is made possible by *opvAB* phase variation, which allows the fast switching of *opvAB* from ON to OFF better states adapted to the new challenges. These results highlight the importance of the reversible switch between ON and OFF expression, as a method for dealing with rapidly varying environments without involving potentially random mutation. Rapidly and specially unpredictably fluctuating environments could select epigenetic gene expression mechanisms for reversible phenotypic plasticity.

## Discussion

Prokaryotes are constantly exposed to changing environmental conditions. Adaptation to these environments is facilitated by prokaryotic biochemical machinery which produces different phenotypic variants. Both mutational and non-mutational mechanisms can contribute to generate this phenotypic variability ^14,35–39^.

In this work, we investigated the adaptative significance of phenotypic heterogeneity in bacterial survival using a well-characterized mechanism that generates phenotypic variation: the epigenetic switch based on the bistable expression of *opvAB* system ^10,11,40^. Here, we set up laboratory evolution experiments to monitor the formation of OpvAB^OFF^ and OpvAB^ON^ lineages in contact with bacteriophages. In alternate cycles of phage presence/absence, *opvAB* bistability persists beyond 50 generations, supporting *opvAB* phase variation as a strategy used by *Salmonella* to survive potential phage encounters. By contrast, continuous phage pressure appears to select for the mutants in genes related to new phage particle assembly and bacterial lysis, as well as genes associated with virulence and cell permeability ^32–34^ (Figs. 1,2). Our results provide clear evidence that in changing environments phenotypic diversity achieved by epigenetic mechanisms is beneficial for the population, whereas, under a constant environment, the maintenance of more than one phenotype appears to be expensive for the population and epigenetic polymorphism is lost, and adapted mutants are selected instead. These findings are in consonance with our mathematical predictions (Fig. 3) and those made by R. Levins in the 60’s ^28,29^ based on game theory, who predicted that genetic polymorphism is advantageous in changing environments but not in stable environments. Therefore, this study extends the Levins predictions made for genetic polymorphism to epigenetic polymorphism^41–45^.

Bacterial adaptation to environmental changes can be facilitated by phenotypic adaptation on a short timescale and by tuning via mutations in the long run ^36^. Both genetic and epigenetic variations are not mutually exclusive and can compete as two strategies of adaptation to fluctuating environments. Using computational approaches, we simulated the epigenetic switching based on *opvAB* system under a range of factors (selection pressure, fluctuation frequency, mutation step-size) to mimic different environmental conditions (Fig. 3). The model simulations seem to adjust quite well to experimental data and predict conditions that favour adaptation by epigenetic switching over mutation depending on the duration and strength of selection pressure. Epigenetic switching is selected over genetic adaptation in fast-changing environments since this strategy seems to minimize adaptation time when the environment fluctuates frequently. These results highlight the importance of epigenetic switching to module the structure of bacterial cell subpopulations in changing environments ^16,17,20,21^.

Microbial populations can evolve a combination of genetic (selection of beneficial mutations) and epigenetic (generation of non-mutational variation that allows adaptation to environmental challenges without mutation) strategies to survive and thrive in dynamic and unpredictable environments. Unlike genetic adaptation, which is a mutational and mostly irreversible mechanism, reversible epigenetic switching provides a high adaptability to adverse agents in natural environments (Fig. 5). Evolution laboratory experiments using multiple selective pressures (exposure to phage, guinea pig serum or macrophages) in cultures containing different proportions of OpvAB^ON^ and OpvAB^OFF^ cells and even cultures with cells blocked in one state (ON or OFF) revealed that only cells that can switch between the OFF and ON states, thanks to the reversibility of *opvAB* phase variation, exhibited the highest survival to added environmental challenges (Fig. 4).These results bring to light the advantages of non-mutational *versus* mutational adaptation mechanisms in unpredictable natural environments (Fig. 5). By switching between phenotypic states, microbial populations can quickly respond to changes in their surroundings, allowing them to take advantage of beneficial conditions and evade unfavourable ones (Fig. 5a). On the contrary, in the case of genetic adaptation, after an environmental change, selection gradually shifts the entire population towards the new optimal phenotype through a mutation or the accumulation of more than one mutation. However, this new phenotype is likely not well-adapted to a new different challenge and the entire bacterial population could be killed (Fig. 5b). Therefore, the possibility of quickly reverting phenotypes provides an added value to epigenetic switching as essential tools for adaptation to changing environments.

**Figure 5.**
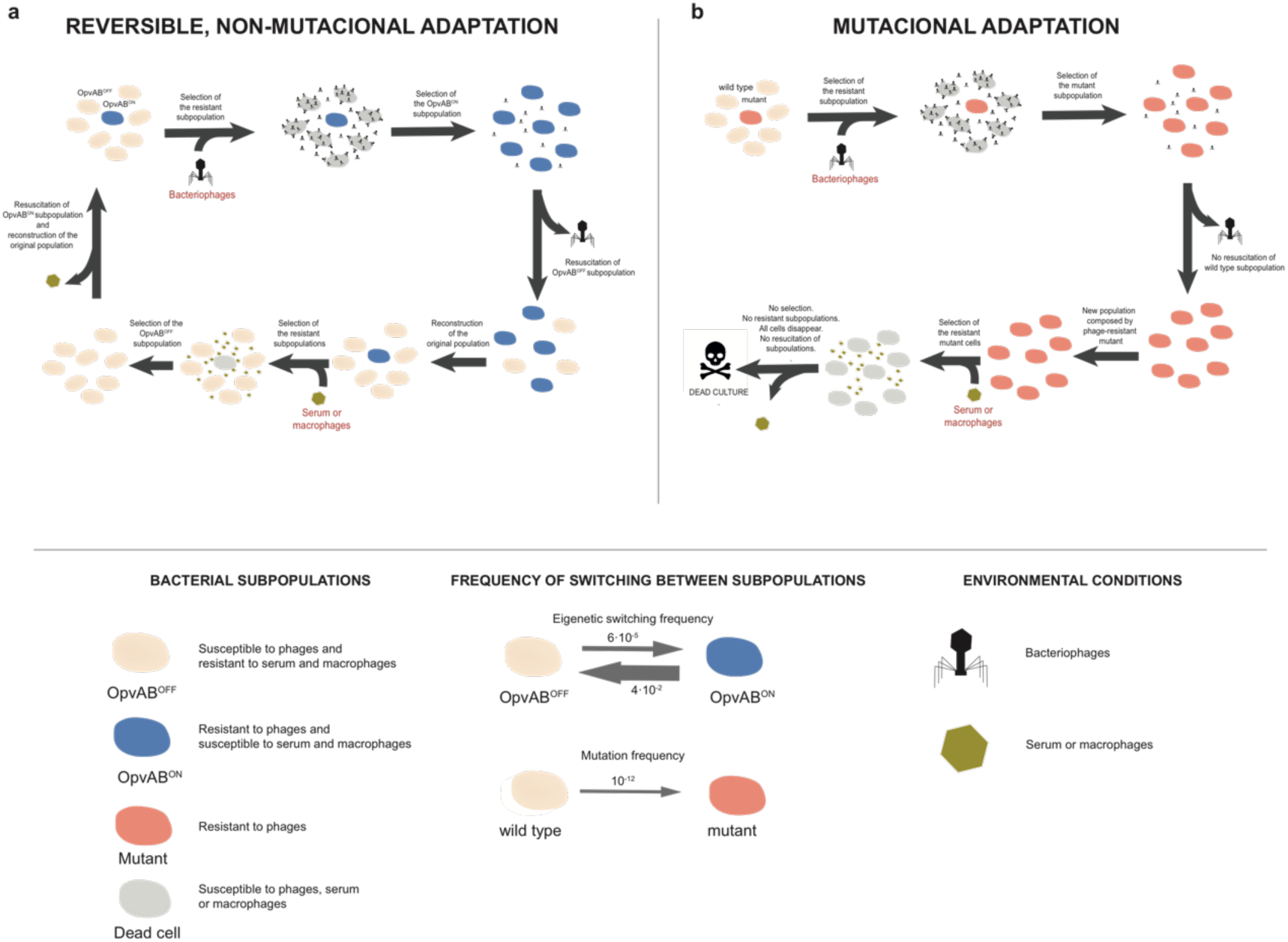
Microbial strategies involved in adaptation to diverse and changing environments controlled by *opvAB* system. Phenotypic variants in a bacterial population can be fashioned by epigenetic (a) and/or genetic mechanisms (b). **a.** Transcription of the *Salmonella enterica opvAB* operon produces lineages of OpvAB^OFF^ and OpvAB^ON^ cells. In the presence of phages, the OpvAB^OFF^ subpopulation will be eliminated, while the OpvAB^ON^ subpopulation will survive. When the phage challenge ceases, OpvAB^OFF^ cells produced by phase variation will survive, and the bistability of OpvAB will be regained. The presence of both OpvAB^OFF^ and OpvAB^ON^ subpopulations will sustain preadaptation to future challenges. In presence of serum or macrophages, the OpvAB^OFF^ subpopulation will be now the survivor, allowing the subsistence of the bacterial culture. **b.** Genetic adaptation involves a beneficial mutation to resist the phage challenge. In the presence of phages, mutant cells will survive and take over the culture; but when phage attacks stop, wild-type cells will not be restored since bistability was lost. The lack of phenotypic switching in culture will reduce the ability of bacterial population to adapt to new and future challenges, where the acquired mutations may not result advantageous. One notable distinction between these strategies is that while epigenetic mechanisms enable reversible switching between different states, genetic adaptation has an irreversible impact in a short time (frequency of switching for both mechanisms is detailed in legend).

This work illustrates the adaptive value of epigenetic variation as a strategy for mutation avoidance in changing and diverse environmental conditions. Phenotypic heterogeneity generated by phase variation may thus allow for enough flexibility to adapt to prompt changes.

## Material and Methods

### Bacterial strains and bacteriophages

*Salmonella enterica* strains listed in Supplementary Table 5 belong to serovar Typhimurium and derive from the mouse-virulent strain ATCC 14028. To simplify, *Salmonella enterica* serovar Typhimurium is often abbreviated as *S. enterica.* Bacteriophages 9NA ^46,47^ was kindly provided by Sherwood Casjens, University of Utah, Salt Lake City.

### Media and growth conditions

Bertani’s lysogeny broth (LB) was used as standard growth medium. Solid media contained agar at 1.5% final concentration. Cultures were grown at 37°C. Aeration of liquid cultures was obtained by shaking at 200 rpm in an Infors Multitron shaker.

### Simulation of fluctuating environments

Presence and absence of bacteriophages were used as conditions to simulate a fluctuating environment (Fig. 1b). The first day of the assay, an overnight culture of *opvAB::lacZ* strain (SV8011) was diluted 1:100 in 5 ml of LB and 100 µl of 9NA lysate (10^9^-10^12^ PFU/ml). After 24 hours of incubation at 37°C with shaking, an aliquot was plated on LB supplemented with X-gal (40 µg/ml) and 100 µl of 9NA bacteriophage. Before the following dilution step, phages were removed using the procedure detailed in the section “Bacteriophage removal procedure”. After phage removal, a dilution was made into 9NA bacteriophage-free LB and incubated for 24 hours at 37°C with shaking. The following day, cells were plated on LB supplemented with X-gal plates (9NA bacteriophage-free). The following dilution and plating were made in presence of 9NA phage, and so on.

Fluctuating conditions were maintained for 7 days: one day adding 9NA phages to cultures and plating on plates with phage, and the following day reducing the number of phages in cultures and plating on phage-free plates. This assay was performed in quadrupled.

### Simulation of constant environments

To simulate a continuous environment (Fig. 1b), an overnight culture of 14028 *opvAB::lacZ* strain (SV8011) was diluted 1:100 in 5 ml of LB with 100 µl 9NA bacteriophage (10^9^-10^12^ PFU/ml). The starting culture was incubated 24 hours at 37°C with shaking at 200 rpm. The following day, an aliquot of this culture was plated on LB supplemented with X-gal plates. These plates had been previously spread with 100 µl of 9NA bacteriophage. Plates were incubated 24 hours at 37°C. These cultures were used for making a new dilution, also in presence of phages. For seven days, dilutions and plating were made in presence of 9NA bacteriophage. This assay was performed in quadrupled.

### Bacteriophage removal procedure

We have fine-tuned a protocol to reduce the number of bacteriophages in cultures. Firstly, to remove bacteriophages in suspension, cultures containing phages were centrifugated for 10 minutes at 4,500 rpm. The supernatant was discarded and bacterial cells from pellet were washed with 10 ml of LB. Cell washing step was repeated three times. The pellet was resuspended in 5 ml of fresh LB. From this bacterial suspension, 50 µl were passed through a sterile 0.45 µm filter using a vacuum filtration system, rinsing the filter with 100 ml of LB. To facilitate phage release from cells, filter containing cells was incubated 2 hours at 37°C without shaking. After this time, the filter was washed with 100 ml of LB. Bacterial cells were recovered from the filter in 2 ml of 2x LB. One hundred µl of this suspension were incubated in the presence of commercial *S. enterica* LPS (Sigma-Aldrich) at a final concentration of 3.75 mg/ml for 3 hours at 37°C with shaking. Twenty µl of this mix were diluted in 1 ml of LB with 0.8 mg/ml of LPS and incubated 2 hours at 37°C with shaking. Hence, phages that are being released into the medium can bind to the commercial LPS instead of bacterial cells. Cells were pelleted by centrifugation 2 minutes at 13,000 rpm and washed three times with LB. Collected cells were resuspended in 20 µl of LB and filtered in a vacuum system using 100 ml of LB. Subsequently, cells were recovered from the filter in 1 ml of 2x LB. Finally, the inoculum for the next day was prepared by adding 100 µl of cells to 1 ml of LB with LPS at 0.45 mg/ml.

### Calculation of number of generations

Number of generations was estimated using the formula X_f_ = X_0_·2^n^ from ^48^. According to this formula, the number of generations (*n*) is calculated as log(X_f_/X_0_)/log(2) where *X_f_* is the number of cells after *n* generations and *X_0_* the initial cell number.

### Isolation of OpvAB^OFF^ phage-resistant mutants

Twenty independent overnight cultures of 14028 *opvAB::lacZ* strain (SV8011) grown in LB were diluted twice in presence of LB with 9NA bacteriophage (10^9^-10^12^ PFU/ml). At least, 10 generations in contact with phage were reached in these cultures.

To isolate colonies, each culture was serially diluted and plated on LB supplemented with X-gal and 100 µl of 9NA lysate agar plates. After incubation at 37°C for 24 h, blue (OpvAB^ON^) and white (OpvAB^OFF^) colonies were visualized.

To ensure that white colonies (OpvAB^OFF^) were not pseudolysogenic, several white colonies were streaked on EBU (Evans Blue Uradine) supplemented with kanamycin agar plates. Pseudolysogenic colonies are visualized as dark-colored colonies. On the contrary, 9NA-free colonies were considered when streaking did not give rise to any dark colony.

### Whole genome DNA sequencing and analysis

Bacterial genome sequencing and bioinformatic analysis were performed by AllGenetics & Biology SL. Genomic DNA was isolated with the Easy-DNA^TM^ kit (Invitrogen), following the manufacturer’s instructions. Quantity and quality of genomic DNA were measured by Qubit High Sensitivity dsDNA Assay (Thermo Fisher Scientific) and Nanodrop ND-1000 spectrophotometer (Thermo Fisher Scientific). Genomic libraries were prepared using the Nextera XT Library Prep kit (Illumina) and its purification was performed by Mag-Bind® RxnPure Plus magnetic beads (Omega Bio-tek). Libraries were pooled in equimolar amounts and sequenced in the platform NovaSeq PE150 run (Illumina), with a total input of 15 gigabases. Sequencing yielded between 3,351,908 and 5,784,360 paired-end reads. The software used to align the reads to their reference genomes was Snippy (https://github.com/tseemann/snippy). SV8011 strain was used as reference genome. Single Nucleotide Polymorphisms (SNPs), insertions and deletions (indels) were localized in *Salmonella* genome by using the *Salmonella enterica* subsp. *enterica* serovar Typhimurium ATCC 14028 genome deposited in GenBank with the accession number CP034230.1.

### Mathematical Modelling

To understand the bistability of *opvAB* genetic system, we considered a model of ON and OFF cell states formation in an environment with continuous exposure to bacteriophage particles and in a fluctuating environment, where phage particles were removed and re-introduced over several intervals. Interactions proposed in Fig. 4a were used to design equations for the rates of change of cell populations in both environments.

The concentration of cells from the initial inoculated strain, referred to as wild-type in Fig. 4a, are represented by *C_off_* and *C_on_* in the equations (Supplementary Table 2). Cells in these concentrations switch between both states at rates *k_off_* and *k_on_* as described in Supplementary Table 3. Since the proportion of cells in the OFF state decreased rapidly after the introduction of bacteriophage particles, this population was considered to be susceptible to infection. This occurred at rate *k_i_* which was also dependent on the concentration of bacteriophage particles, represented by *V*. Because time passes between the infection of a cell by a phage particle and the release of new particles by bursting, the infected cells were split into two states (early infected cells and late infected cells). The concentration of cells early after infection, represented by *C_E_*, increases as cells are infected and continue to replicate before transitioning at rate *k_1_* to become late infected cells. The concentration of late infected cells is represented by *C_L_*. This late infected cell concentration decreases as cells burst at rate *k_2_* releasing *b* new phage particles.

In the constant bacteriophage environment, the proportion of cells in the OFF state recovered gradually after the initial decrease following the introduction of the particles. To model this, wild-type cells became resistant mutants at rate *k_m_*, which continue to undergo switching at rates *k_off_* and *k_on_.* The concentrations of the mutant cells in the two states are represented by *C_m,off_* and *C_m,on_*. As no additional selective pressures on the mutated populations were introduced, this transition was considered permanent for the model.

Phage particles were assumed to be stable in the growth media, so no decay term was included in equation (8).

The proportion of cells in the OFF state expected to be observed using the plate screening technique described in the experimental procedure was calculated using equation (9). Infected cells were assumed to not form visible colonies.

To simplify the calculations, the maximum concentration of cells that can be supported by the growth media was normalized to 1. The experimental workflow describes using a 1:100 dilution of cells to start the constant environment, so the total concentration of initial cells was estimated in equation (10).

From previous experiments, starting proportions of wild-type cells were assumed to be approximately 0.998 in the OFF state and 0.002 in the ON state, so the initial cell concentration was split into these two states (equation 11 and 12).

Because the cells were taken from incubation of a single strain and with no bacteriophage particles in the environment, initial mutant and infected cell concentrations were assumed to be 0 (equation 13).

The number of phage particles introduced at the beginning was approximated to be 122 per cell, based on the experimental description. The initial concentration of cells was used to calculate the initial concentration of bacteriophage particles (equation 14).

Some rates were rewritten in terms of others to reduce the number of parameters varied for fitting. From the observed proportions of OFF and ON states, *k_on_* was estimated to be close to 0.002 times *k_off_* (equation 15).

From experimental assays, total lysis time using 122 bacteriophage particles per cell was approximately 4 hours. This was converted to generations using the estimated generation time of 43 minutes (equation 16)

The total time to lyse for a cell in the model is the sum of the inverse of *k_1_* and *k_2_*, as shown in equation (17), so *k_2_* was defined in terms of *k_1_* and the total lysis time in generations.

Because the data was available as proportion of cells in OFF state, the predicted proportion of OFF cells from the model was calculated from the concentrations using equation (9). The fitting was performed by minimizing the least squares error between the available data and *P_off_* predicted proportion by the model. For this minimization, parameters *k_off_*, *k_m_*, *k_i_*, *k_1_*, and *b* were varied, from which *k_on_* and *k_2_*, were calculated using equations (15) and (17). Fitting was performed in Excel using the Solver tool with the GRG Nonlinear method. Table Supplementary Table 4 showed the best-fit parameters results from constant bacteriophage environment.

For the fluctuating environment case, the same population dynamic equations (1) through (8) were used and the proportion of OFF cells calculated using equation (9) was used for plotting. The best fit rates that were determined in the constant environment in Supplementary Table 4 were expected to be good estimates for the switching, mutation, and infection interactions, and so these were applied in equations (1) through (8) to calculate the populations in the fluctuating environment.

Since the first interval included phages, the calculations were started with the same initial conditions as the constant environment shown in equations (10) through (14). At the end of this interval, bacteriophage particles were removed to create a phage-free environment by a cleaning process. The model calculations were stopped and then solved using new initial conditions at this point.

From the experimental procedure description, a 1:100 dilution factor was used to calculate the new initial concentrations for the wild-type populations (equation 18).

In the fluctuating environment, the proportion of OFF cells recovered after bacteriophage particles were removed, but then decreased when bacteriophage particles were reintroduced. Therefore, it was assumed that the proportion of phage-resistant mutants was small and subject to extinction at dilution events. The concentrations were reset to 0 for this (equation 19).

Bacteriophage particles, early infected, and late infected cells were assumed to be completely removed in the cleaning process, so these concentrations were also reset to 0 (equation 20).

At the beginning of intervals when bacteriophage particles were re-introduced, the model was again stopped, then solved with new initial conditions. The cells were transferred to the new environment using the same 1:100 dilution, so the same new initial condition equations (18) and (19) were used. Early and late infected cell concentrations were still expected to be 0 as in equations since there were no bacteriophage particles in the preceding interval. Since the same experimental set up was used, the concentration of introduced bacteriophage particles was reused from equation (14).

### Measurement of survival in guinea-pig serum

Survival in guinea pig serum (Sigma-Aldrich) was analyzed as described in ^49^ with some modifications. Overnight cultures of *S. enterica* were diluted 1:100 in LB and incubated for 2 hours at 37°C with shaking (200 rpm) until O.D._600 nm_ ∼ 0.5 (3·10^8^ CFU/ml). Bacterial cells were pelleted by centrifugation at 4,500 rpm 20 minutes and resuspended in fresh LB medium. Cells were serially diluted in PBS supplemented with 2 mM MgCl_2_ in order to reach 2·10^4^ CFU/ml. Guinea pig serum was added to 30% (v/v) final concentration and mixtures were incubated at 37°C without shaking. Samples were taken every 30 minutes by plating on LB supplemented with X-gal agar plates. Viable counts were expressed as a percentage of initial concentration (% survival). Each strain was assayed in triplicate. The ordinary one-way ANOVA was used to determine if the differences in survival to serum observed in different backgrounds were statistically significant.

To analyse the dynamic proportions of ON and OFF states, we utilized overnight cultures of *S. enterica* that were cultivated with 9NA bacteriophage for 10 generations. After exposure in guinea pig serum at different times, we determined the ratio of OpvAB^ON^ and OpvAB^OFF^ colonies by counting them in LB plates supplemented with X-gal incubated at 37°C for 24 hours.

### Macrophage proliferation assay

The rate of intramacrophage proliferation after 18 hours of infection was determined using J774 murine macrophages cell line ^50^. The day before infection, J774 cells were plated in 24-well plates (Thermo Scientific) at 2.5·10^5^ cells/well density in DMEM supplemented with 10% (v/v) fetal bovine serum (Biowest). Two 24-well plates were used: one was used to monitor internalized bacteria after 30 minutes of infection (30min_plate), while the other one was utilized to determine the bacterial proliferation inside macrophages after 18 hours of the infection (18h_plate). Plates were incubated 24 hours at 37°C with 5% CO_2_.

To perform the macrophage cell infection, *S. enterica* overnight cultures were collected by centrifugation, washed, resuspended in LB and diluted in DMEM to the required concentration. Bacteria were added to the wells using a multiplicity of infection (MOI) of 10 bacteria per macrophage and incubated at 37°C with 5% CO_2_. Thirty minutes after infection, cells were washed twice with sterile PBS 1x and incubated with fresh DMEM supplemented with 100 µg/ml gentamicin for 90 minutes. After 90 minutes of incubation, cells were washed twice with PBS. While macrophages from 30min_plate were lysed, macrophages from 18h_plate were incubated with DMEM supplemented with gentamicin at 16 µg/ml and incubated for 18 additional hours before lysis. The number of CFU per well was calculated after incubation with 1% Triton X-100 (Sigma-Aldrich) in PBS for 10 minutes at 37°C to release bacteria, plating appropriate dilutions on LB agar and counting colonies after 24 hours of incubation at 37°C.

The intramacrophage proliferation rate was determined as the ratio between the number of bacteria at 18 hours and the number of internalized bacteria after 30 minutes phagocytosis. Infections were carried out in triplicate. The ordinary one-way ANOVA was used to determine if the differences in replication rates observed in different backgrounds were statistically significant.

For the dynamic analysis of ON and OFF proportions, *S. enterica* overnight cultures grown in presence of 9NA bacteriophage for 10 generations were used. Proportions of OpvAB^ON^ and OpvAB^OFF^ colonies were calculated after counting colonies after 24 hours of incubation at 37°C in plates containing LB supplemented with X-gal.

### Statistical analysis

Statistical analyses were performed using GraphPad Prism version 9.0.0. All experiments were performed at least three times. One-way ANOVA was performed for multiple comparisons. *p* value < 0.05 was considered statistically significant.

## Acknowledgments

This study was supported by grants BIO2016-75235 from the Ministerio de Ciencia e Innovación of Spain, the grant PID2020-116995RB-I00 funded by MCIN/AEI/ 10.13039/5011100011033 and, as appropriate, by “ERDF A way of making Europe”, by the “European Union” or by the “European Union Next Generation EU/PRTR”, the grant R35GM148351 from NIH-NIGMS and the VI Plan Propio de Investigación y Transferencia from the Universidad de Sevilla. We are grateful to P. Bernal, C. Beuzón, M. Olmedo, E. Mellado-Durán and L. Bossi for critically reading the manuscript and for helpful comments and suggestions.

## SUPPORTING INFORMATION

**Supplementary Fig. 1.** Equations for explicating of the bistable *opvAB* system mathematical modelling.

**Supplementary Fig. 2.** Version of constant environment fit with experimental data. Grey dots represent the experimental data from the four independent experiments graphed in Fig. 1d.

**Supplementary Table 1.** Characterization of 20 OpvAB^OFF^ phage-resistant mutant isolates. Affected genes, predicted functions, abundancy in isolates and biological process involved.

**Supplementary Table 2.** Variable terms used in model equations for cell and bacteriophage populations.

**Supplementary Table 3.** Rate parameters for model equations.

**Supplementary Table 4.** Best-fit parameter results from constant bacteriophage environment.

**Supplementary Table 5.** List of strains used in this work.

