## Supplementary Fig. 1 for "Evolution of a bistable genetic system in fluctuating and non-fluctuating environments"

Model equations

$$C_{tot} = C_{off} + C_{on} + C_{m,off} + C_{m,on} + C_E + C_L \qquad (1)$$

$$\frac{\mathrm{d}C_{off}}{\mathrm{dt}} = \underbrace{C_{off} \cdot (1 - C_{tot})}_{\text{Replication}} + \underbrace{K_{off} \cdot C_{on}}_{\substack{\text{Switch from} \\ \text{wt ON}}} - \underbrace{K_{on} \cdot C_{off}}_{\substack{\text{Switch from} \\ \text{wt ON}}} - \underbrace{K_m \cdot C_{off}}_{\text{Mutation}} - \underbrace{K_i \cdot C_{off} \cdot V}_{\text{Infection}} \qquad (2)$$

$$\frac{\mathrm{d}C_{on}}{\mathrm{dt}} = \underbrace{C_{on} \cdot (1 - C_{tot})}_{\text{Replication}} - \underbrace{K_{off} \cdot C_{on}}_{\substack{\text{Switch from} \\ \text{wt OFF}}} + \underbrace{K_{on} \cdot C_{off}}_{\substack{\text{Switch from} \\ \text{wt OFF}}} - \underbrace{K_m \cdot C_{on}}_{\text{Mutation}} \qquad (3)$$

$$\frac{\mathrm{d}C_{m,off}}{\mathrm{dt}} = \underbrace{C_{m,off} \cdot (1 - C_{tot})}_{\text{Replication}} + \underbrace{K_{off} \cdot C_{m,on}}_{\substack{\text{Switch from} \\ \text{mutant ON}}} - \underbrace{K_{on} \cdot C_{m,off}}_{\substack{\text{Switch to} \\ \text{mutant ON}}} + \underbrace{K_m \cdot C_{off}}_{\text{Mutation}} \qquad (4)$$

$$\frac{\mathrm{d}C_{m,on}}{\mathrm{dt}} = \underbrace{C_{m,on} \cdot (1 - C_{tot})}_{\text{Replication}} - \underbrace{K_{off} \cdot C_{m,on}}_{\substack{\text{Switch to} \\ \text{mutant OFF}}} + \underbrace{K_{on} \cdot C_{m,off}}_{\substack{\text{Switch from} \\ \text{mutant OFF}}} + \underbrace{K_m \cdot C_{on}}_{\text{Mutation}} \qquad (5)$$

$$\frac{\mathrm{d}C_E}{\mathrm{dt}} = \underbrace{C_E \cdot (1 - C_{tot})}_{\text{Replication}} + \underbrace{K_i \cdot C_{off} \cdot V}_{\text{Infection}} - \underbrace{K_1 \cdot C_E}_{\substack{\text{Transition to} \\ \text{late infected}}} \qquad (6)$$

$$\frac{\mathrm{d}C_L}{\mathrm{dt}} = \underbrace{K_1 \cdot C_E}_{\substack{\text{Transition} \\ \text{from early} \\ \text{infected}}} - \underbrace{K_2 \cdot C_L}_{\text{Bursting}} \qquad (7)$$

$$\frac{\mathrm{dV}}{\mathrm{dt}} = \underbrace{b \cdot K_2 \cdot C_L}_{\substack{\text{Release from} \\ \text{bursting}}} - \underbrace{K_i \cdot C_{off} \cdot V}_{\text{Infection}} \qquad (8)$$

$$P_{off} = \frac{C_{off} + C_{m,off}}{C_{off} + C_{m,off} + C_{on} + C_{m,on}} \qquad (9)$$

$$C_{tot}(0) = 1 \cdot \left(\frac{1}{100}\right) dilution = 0.01 \qquad (10)$$

$$C_{off}(0) = 0.998 \cdot C_{tot}(0) = 0.00998 \qquad (11)$$

$$C_{on}(0) = 0.002 \cdot C_{tot}(0) = 0.00002 \qquad (12)$$

$$C_{m,off}(0) = C_{m,on}(0) = C_E(0) = C_L(0) = 0 \qquad (13)$$

$$\mathrm{V}(0) = 122\ particles\ per\ cell \cdot C_{tot}(0)\ cells = 1.22\ \text{particles} \qquad (14)$$

$$K_{on} = 0.002 \cdot K_{off} \qquad (15)$$

$$\text{Total lysis time} = 4\ hours \cdot \frac{60\ minutes}{1\ hour} \cdot \frac{1\ generation}{43\ minutes} = 5.6\ generations \qquad (16)$$

$$\text{Total lysis time} = \frac{1}{K_1} + \frac{1}{K_2} \rightarrow K_2 = \left(5.6 - \frac{1}{K_1}\right)^{-1} \qquad (17)$$

$$C_{off}(0) = \frac{1}{100} \cdot C_{off}(t), C_{on}(0) = \frac{1}{100} \cdot C_{on}(t) \qquad (18)$$

$$C_{m,off}(0) = C_{m,on}(0) = 0 \qquad (19)$$

$$C_E(0) = C_L(0) = V(0) = 0 \qquad (20)$$
