## Supplementary figures and images for "Evolution of a bistable genetic system in fluctuating and non-fluctuating environments"

### Supplementary Fig. 2

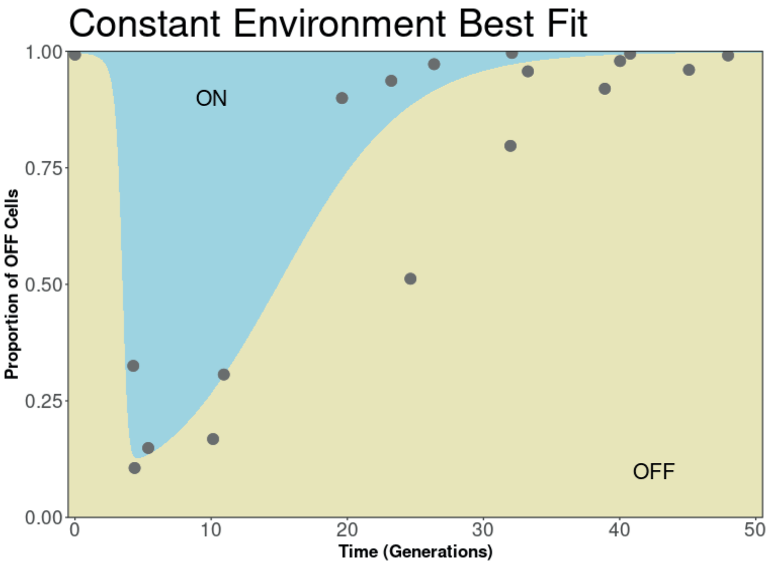
