## Supplementary Tables for "Evolution of a bistable genetic system in fluctuating and non-fluctuating environments"

### SUPPORTING INFORMATION

**Supplementary Table 1.** Characterization of 20 OpvAB$^{OFF}$ phage-resistant mutant isolates. Affected genes, predicted functions, abundancy in isolates and biological process involved.

| Gene ID | Gene name | Gene function/activity | % of mutant isolates containing mutations | Biological process involved |
|---|---|---|---|---|
| EJJ31_RS13680 | | S49 family peptidase/Capsid maduration | 90 | Phage infection and bacterial lysis |
| EJJ31_22835 | *tuf* | Traslational elongation factor Tu | 85 | Translation, ribosomal structure and biogenesis |
| EJJ31_RS13670 | | Major capsid protein/Stabilization of the condensed DNA in phage heads | 70 | Phage infection and bacterial lysis |
| EJJ31_RS13580 | | Phage virulence factor PagK family protein secreted by SPI-2 T3SS | 65 | Virulence |
| EJJ31_15335 | *gtgA* | Type III secretion system effector protease GtgA | 60 | Virulence |
| EJJ31_RS24715 | | IS3 family transposase | 60 | Mobilome: prophages, transposons |
| EJJ31_RS13650 | | RNA-guided endonuclease TnpB family protein/Functional progenitor of CRISPR-Cas nucleases | 60 | Defense mechanisms |
| EJJ31_RS24730 | | DUF1983 domain-containing protein/host specificity protein J | 55 | Phage infection and bacterial lysis |
| EJJ31_RS13700 | | Terminase small subunit/DNA packaging enzyme of bacteriophages | 55 | Phage infection and bacterial lysis |
| EJJ31_RS06080 | | PPC domain-containing protein | 55 | Unknown |
| EJJ31_RS14845 | | Minor phage tail protein | 55 | Phage infection and bacterial lysis |

| | | | | |
|---|---|---|---|---|
| EJJ31_RS15020 | | DinI family protein | 50 | SOS response |
| EJJ31_RS13710 | | Lysis protein involved in disruption of the outer membrane by bacteriophages | 50 | Phage infection and bacterial lysis |
| EJJ31_RS10810 | | rtT sRNA, processed from tyrT transcript | 50 | Translation, ribosomal structure and biogenesis |
| EJJ31_RS13675 | | Phage head decoration protein | 45 | Phage infection and bacterial lysis |
| EJJ31_RS16130 | | Oxaloacetate decarboxylase subunit gamma/Sodium ion export | 45 | Inorganic ion transport and metabolism |
| EJJ31_RS04145 | | Nickel/cobalt efflux protein RcnA | 40 | Inorganic ion transport and metabolism |
| EJJ31_06200 | *gogA* | Type III secretion system effector protease GogA | 40 | Virulence |
| EJJ31_RS14840 | | Tail fiber protein required for assembly of the distal part of the long fibres | 35 | Phage infection and bacterial lysis |
| EJJ31_RS13665 | | Protein FI required for efficient phage DNA packaging and in head assembly | 30 | Phage infection and bacterial lysis |
| EJJ31_RS09365 | | DUF968 domain-containing protein that acts as recombinase that participates in DNA repair and replication | 25 | Replication, recombination and repair |
| EJJ31_RS22390 | | rRNA-16S ribosomal RNA | 25 | Translation, ribosomal structure and biogenesis |
| EJJ31_RS23095 | | rRNA-23S ribosomal RNA | 25 | Translation, ribosomal structure and biogenesis |
| EJJ31_RS18545 | | DUF6531 domain-containing protein | 20 | Unknown |
| EJJ31_RS14960 | | Lysozyme required for host cell lysis | 20 | Phage infection and bacterial lysis |

| | | | | |
|---|---|---|---|---|
| EJJ31_RS02500 | | Oxalacetate decarboxylase subunit beta involved in sodium ion transport | 20 | Inorganic ion transport and metabolism |
| EJJ31_04980 | *sipD* | SPI-1 type III secretion system needle tip complex protein SipD | 20 | Virulence |
| EJJ31_15195 | *sseI* | SPI-2 type III secretion system effector SseI | 20 | Virulence |
| EJJ31_RS05470 | | FliC/FljB family flagellin | 15 | Motility |
| EJJ31_RS03125 | | TerC integral membrane family proteins related to tellurium ions efflux | 15 | Inorganic ion transport and metabolism |
| EJJ31_RS13590 | | Prophage tail fiber N-terminal domain-containing protein | 15 | Phage infection and bacterial lysis |
| EJJ31_RS10190 | | Tyrosine-type recombinase/integrase | 15 | Replication, recombination and repair |
| EJJ31_RS13660 | | Phage head-tail joining protein | 10 | Phage infection and bacterial lysis |
| EJJ31_RS13600 | | Host specificity protein J that attaches the phage to the host receptor, inducing viral DNA injection | 10 | Phage infection and bacterial lysis |
| EJJ31_RS14890 | | Phage tail tape measure protein, important for assembly of phage tails and involved in tail length determination | 10 | Phage infection and bacterial lysis |
| EJJ31_02310 | *rrf* | rRNA-5S ribosomal RNA | 10 | Translation, ribosomal structure and biogenesis |
| EJJ31_20180 | *oadA* | Sodium-extruding oxaloacetate decarboxylase subunit alpha/sodium ion transport | 10 | Inorganic ion transport and metabolism |
| EJJ31_RS13820 | | CII family transcriptional regulator | 10 | Transcription |
| EJJ31_08180 | *sspH2* | SPI-2 type III secretion system effector E3 ubiquitin transferase SspH2 | 5 | Virulence |

| | | | | |
|---|---|---|---|---|
| EJJ31_13020 | *sseA* | SPI-2 type III secretion system chaperone SseA | 5 | Virulence |
| EJJ31_05070 | *avrA* | Type III secretion system YopJ family effector AvrA | 5 | Virulence |
| EJJ31_RS24685 | | DNA breaking-rejoining protein with nucleic acid phosphodiester bond hydrolysis activity | 5 | Replication, recombination and repair |
| EJJ31_06755 | *shdA* | Large outer membrane protein fibronectin-binding adhesin ShdA | 5 | Cell wall/membrane/envelope biogenesis |
| EJJ31_RS13565 | | IS481 family transposase | 5 | Mobilome: prophages, transposons |
| EJJ31_RS13810 | | Replication protein 14 | 5 | Replication, recombination and repair |
| EJJ31_15900 | *rlmC* | 23S rRNA (uracil(747)-C(5))-methyltransferase RlmC | 5 | Defense mechanisms |
| EJJ31_RS03425 | | Anion permease | 5 | Inorganic ion transport and metabolism |
| EJJ31_RS23145 | | Aryl-sulfate sulfotransferase | 5 | Unknown |
| EJJ31_01885 | *bigA* | Autotransporter adhesin BigA | 5 | Cell wall/membrane/envelope biogenesis |
| EJJ31_15155 | *zapC* | Cell division protein ZapC | 5 | Cell cycle control, cell division and chromosome partitioning |
| EJJ31_RS15675 | | Metal ion binding CoA ester lyase | 5 | Inorganic ion transport and metabolism |
| EJJ31_RS23600 | | FadR family transcriptional regulator | 5 | Transcription |
| EJJ31_RS19025 | | Fimbrial biogenesis outer membrane usher protein | 5 | Motility |
| EJJ31_RS19120 | | Fimbrial protein | 5 | Motility |
| EJJ31_RS19935 | | Glycosyl hydrolase family 18 protein | 5 | Carbohydrate transport and metabolism |
| EJJ31_22105 | *lpxO* | Lipid A hydroxylase LpxO | 5 | Cell wall/membrane/envelope biogenesis |

| | | | | |
|---|---|---|---|---|
| EJJ31_RS16380 | | Membrane protein | 5 | Cell wall/membrane/envelope biogenesis |
| EJJ31_RS03560 | | Methyl-accepting chemotaxis protein | 5 | Signal transduction mechanisms |
| EJJ31_00675 | *rfaL* | O-antigen ligase RfaL | 5 | Cell wall/membrane/envelope biogenesis |
| EJJ31_RS06295 | | OFA family MFS transporter | 5 | Carbohydrate transport and metabolism |
| EJJ31_RS12800 | | PfkB family carbohydrate kinase with DNA-binding transcription factor activity | 5 | Transcription |
| EJJ31_RS13685 | | Phage portal protein | 5 | Phage infection and bacterial lysis |
| EJJ31_10355 | *edd* | Phosphogluconate dehydratase | 5 | Carbohydrate transport and metabolism |
| EJJ31_RS19115 | | PTS sugar transporter subunit IIA | 5 | Carbohydrate transport and metabolism |
| EJJ31_RS03175 | | Siderophore-interacting protein involved in iron transport | 5 | Inorganic ion transport and metabolism |
| EJJ31_RS17115 | | Sigma-54-dependent transcriptional regulator | 5 | Transcription |
| EJJ31_08535 | *dusC* | tRNA dihydrouridine(16) synthase DusC | 5 | Translation, ribosomal structure and biogenesis |
| EJJ31_RS12595 | | tRNA-Val | 5 | Translation, ribosomal structure and biogenesis |
| EJJ31_17295 | *uspG* | Universal stress protein UspG | 5 | SOS response |
| EJJ31_RS16720 | | UxaA family hydrolase involved in D-galacturonate catabolic process | 5 | Carbohydrate transport and metabolism |
| EJJ31_RS21485 | | YjiK family protein | 5 | Unknown |
| EJJ31_RS11970 | | Zinc-binding alcohol dehydrogenase family protein with oxidoreductase activity | 5 | Energy production and conversion |

**Supplementary Table 2.** Variable terms used in model equations for cell and bacteriophage populations

| **Population** | **Definition** |
|---|---|
| $C_{off}$ | Concentration of wild-type cells with operon in OFF state |
| $C_{on}$ | Concentration of wild-type cells with operon in ON state |
| $C_{m,off}$ | Concentration of mutant cells with operon in OFF state |
| $C_{m,on}$ | Concentration of mutant cells with operon in ON state |
| $C_E$ | Concentration of early infected cells that replicate and do not yet lyse |
| $C_L$ | Concentration of late infected cells that do not replicate and actively lyse |
| $C_{tot}$ | Total concentration of all cells in media |
| $V$ | Concentration of bacteriophage particles |

**Supplementary Table 3.** Rate parameters for model equations

| **Rate** | **Definition** |
|---|---|
| $k_{off}$ | Rate of cells in ON state switching to OFF state |
| $k_{on}$ | Rate of cells in OFF state switching to ON state |
| $k_m$ | Rate of wild-type cells becoming resistant mutants |
| $k_i$ | Rate of infection of susceptible cells |
| $k_1$ | Rate of early infected cells transitioning to late infected state |
| $k_2$ | Rate of late infected cells bursting |
| $b$ | Number of bacteriophage particles released by a late infected cell |

**Supplementary Table 4.** Best-fit parameter results from constant bacteriophage environment

| **Parameter** | **Fitted result** |
|---|---|
| $k_{off}$ | 0.21 per generation per cell |
| $k_{on}$ | $4.2 \cdot 10^{-4}$ per generation per cell, calculated from equation 15 |
| $k_m$ | $9.55 \cdot 10^{-5}$ per generation per cell |
| $k_i$ | 0.667 per generation per cell |
| $k_1$ | 1.05 per generation per cell |
| $k_2$ | 0.215 per generation per cell, calculated from equation 17 |
| *b* | 303 phage particles per cell |

**Supplementary Table 5.** List of strains used in this work.

| **Strain** | **Genotype** |
|---|---|
| ATCC 14028 | Wild type |
| SV8011[a] | 14028 *opvAB::lacZ* |
| SV6401[a] | 14028 $\text{OpvAB}^{\text{ON}}$ |
| SV8020[a] | 14028 Δ*opvAB LPS mutant* |
| SV6796[a] | 14028 $\Delta wzz_{ST}$ $\Delta wzz_{fepE}$ |
| SV6063 | 14028 Δ*SPI-2* |
| SV10143 | SV8011 Mutant 1 |
| SV10144 | SV8011 Mutant 2 |
| SV10145 | SV8011 Mutant 3 |
| SV10146 | SV8011 Mutant 4 |
| SV10147 | SV8011 Mutant 5 |
| SV10148 | SV8011 Mutant 6 |
| SV10149 | SV8011 Mutant 7 |
| SV10150 | SV8011 Mutant 8 |
| SV10151 | SV8011 Mutant 9 |
| SV10152 | SV8011 Mutant 10 |
| SV10153 | SV8011 Mutant 11 |
| SV10154 | SV8011 Mutant 12 |

| | |
|---|---|
| SV10155 | SV8011 Mutant 13 |
| SV10156 | SV8011 Mutant 14 |
| SV10157 | SV8011 Mutant 15 |
| SV10158 | SV8011 Mutant 16 |
| SV10159 | SV8011 Mutant 17 |
| SV10160 | SV8011 Mutant 18 |
| SV10161 | SV8011 Mutant 19 |
| SV10162 | SV8011 Mutant 20 |

[a] Strains described in [10].
